# Cell-intrinsic ceramides determine T cell function during melanoma progression

**DOI:** 10.1101/2022.08.31.505996

**Authors:** Matthias Hose, Anne Günther, Eyad Naser, Fabian Schumacher, Tina Schönberger, Julia Falkenstein, Athanassios Papadamakis, Burkhard Kleuser, Katrin Anne Becker, Erich Gulbins, Adriana Haimovitz-Friedman, Jan Buer, Astrid M. Westendorf, Wiebke Hansen

**Author notes:** Corresponding author: Prof. Dr. Wiebke Hansen, Institute of Medical Microbiology, University Hospital Essen, University Duisburg-Essen, Hufelandstr. 55, 45147 Essen, Germany. contributed equally (Matthias Hose, Anne Günther).

## Abstract

Acid sphingomyelinase (Asm) and acid ceramidase (Ac) are parts of the sphingolipid metabolism. Asm hydrolyzes sphingomyelin to ceramide, which is further metabolized to sphingosine by Ac. Ceramide generates ceramide-enriched platforms that are involved in receptor clustering within cellular membranes. However, the impact of cell-intrinsic ceramide on T cell function is not well characterized. By using T cell-specific Asm- or Ac-deficient mice, with reduced or elevated ceramide levels in T cells, we identified ceramide to play a crucial role in T cell function *in vitro* and *in vivo*. T cell-specific ablation of Asm in Asm^flox/flox^/CD4cre mice resulted in enhanced tumor progression associated with impaired T cell responses, whereas Ac^flox/flox^/CD4cre mice showed reduced tumor growth rates and elevated T cell activation compared to the respective controls upon tumor transplantation. Further *in vitro* analysis revealed that decreased ceramide content supports CD4^+^ regulatory T cell differentiation and interferes with cytotoxic activity of CD8^+^ T cells. In contrast, elevated ceramide concentration in CD8^+^ T cells from Ac^flox/flox^/CD4cre mice was associated with enhanced cytotoxic activity. Strikingly, ceramide co-localized with the T cell receptor (TCR) and CD3 in the membrane of stimulated T cells and phosphorylation of TCR signaling molecules was elevated in Ac-deficient T cells. Hence, our results indicate that modulation of ceramide levels, by interfering with the Asm or Ac activity has an effect on T cell differentiation and function and might therefore represent a novel therapeutic strategy for the treatment of T cell-dependent diseases such as tumorigenesis.

## Introduction

Sphingolipids are structural components and bioactive molecules of cellular membranes with different functions in cellular processes. One important member of the sphingolipid family is ceramide. Ceramide has the ability to form ceramide-enriched platforms within the plasma membrane [1]. These microdomains contribute to receptor clustering and other protein interactions and are thereby involved in several signaling pathways and important cellular processes including proliferation, migration, differentiation, and apoptosis [2].

The enzyme acid sphingomyelinase (Asm) generates ceramide by hydrolyzing sphingomyelin, whereas acid ceramidase (Ac) converts ceramide into sphingosine [3]. Dysregulations in enzyme activity of the ceramide metabolism can lead to severe diseases. Specifically, mutations in the *SMPD1* gene, encoding for Asm, resulting in loss of or reduced Asm activity, cause the lipid-storage disease Niemann-Pick type A and B (NPD) [4]. NPD type A patients suffer from severe neuronal symptoms, due to sphingomyelin accumulations in the central nervous system [5]. In contrast, elevated Asm activity is associated with the development of major depressive disorders [6]. Pharmacological inhibitors of Asm (functional inhibitors of acid sphingomyelinase, FIASMA) are therefore used as antidepressant drugs [7]. Loss of Ac activity leads to the development of Farber Disease (FD). FD patients suffer from arthralgia, hepatosplenomegaly and a general developmental delay [8].

In addition, alterations in the sphingolipid metabolism play an important role in other pathological disorders including different tumor entities [9]. For instance, Asm-generated ceramide-enriched platforms are crucial for the CD95-mediated induction of apoptosis [10]. Cancer cells may upregulate Ac expression, leading to reduced ceramide abundance and increased levels of pro-survival lipid spingosine-1-phosphate (S1P) and thereby foster their survival [11]–[13]. Therefore, targeting of Asm and Ac in experimental cancer therapies have shown anti-tumoral efficacy. For example, ionizing radiation induces the activity of Asm, leading to ceramide generation and apoptosis of cancer cells, but fails to do so in lymphoblasts from NPD patients, which lack Asm activity [14]. In addition, Mauhin et. al. demonstrated most recently, in a retrospective study, that there was an elevated incidence for cancer in NPD patients [15]. We previously observed an enhanced tumor growth rate of transplanted tumor cells in Asm-deficient mice as compared to wildtype (WT) mice. Investigation of the tumor microvasculature identified apoptosis-resistant endothelial cells in Asm-deficient mice as a driver of elevated tumor growth [16]. In various cancer cell lines Ac facilitates proliferation by the degradation of ceramide [17]. Pharmacological inhibition of Ac has been described to be effective for the treatment of patients suffering from colorectal cancer [18].

Emphasizing the important role of ceramide in the modulation of tumorigenesis, recent studies by Ghosh et. al. provided evidence that the application of exogenous C2 ceramide induces a strong anti-tumor response by increasing frequencies of cytotoxic CD8^+^ and IFN-γ-producing CD4^+^ T cells [19]. Among other immune cells, T cells play a crucial role during tumor progression. In melanoma patients the ratio of cytotoxic CD8^+^ T cells versus CD4^+^Foxp3^+^ regulatory T cells (Tregs) in the tumor microenvironment is predictive for the disease outcome [20]. The infiltration of Tregs into the tumor tissue is considered as a critical step during tumorigenesis. We provided evidence that Neuropilin-1 (Nrp-1), highly expressed by Tregs [21], regulates the migration of Tregs into VEGF-producing tumor tissue accompanied by elevated tumor progression [22]. Depletion of Tregs improved the anti-tumoral immune response of CD8^+^ T cells in colitis-associated colon cancer, emphasizing the important role of the T cell composition in tumorigenesis [23].

Analysis of immune responses in Asm-deficient mice have shown an impaired cytotoxic activity of CD8^+^ T cells during LCMV infection [24]. Moreover, Asm has been identified as a negative regulator of Treg development [25]. Asm-deficient or FIASMA-treated mice showed increased numbers of Tregs in comparison to WT animals [26]. In accordance, we detected lower Treg frequencies in spleens of t-Asm/CD4cre mice, which overexpress Asm specifically in T cells [27]. In CD4^+^ T cells, isolated from human peripheral blood, inhibition of Asm led to an impaired T cell receptor (TCR) signal transduction accompanied by reduced T cell proliferation and impaired CD4^+^ T helper (Th) cell differentiation [28]. The role of Ac in T cell responses is largely unclear. Nevertheless, as mentioned above, several lines of evidence indicate that ceramide metabolism participates in the regulation of T cell responses.

Here, we demonstrate that elevated ceramide concentrations facilitate TCR signaling cascades and determine T cell activation and differentiation *in vitro*. Strikingly, transplantation of B16-F1 melanoma cells into Asm^flox/flox^/CD4cre or Ac^flox/flox^/CD4cre mice, which exhibit decreased or increased ceramide levels in T cells respectively, revealed that increased ceramide concentrations improve the anti-tumoral T cell response during melanoma progression.

## Materials and Methods

### Mice

All mice were on C57BL/6 background and maintained under specific pathogen-free conditions at the Animal Facility of University Hospital Essen. Female C57BL/6 mice were purchased from Envigo Laboratories (Envigo CRS GmbH, Rossdorf, Germany). Asm-KO (Smpd1^tm1Esc^) mice has been described previously [29]. Asm^flox/flox^ (Smpd1^tm1a(EUCOMM)Wtsi^) mice and Ac^flox/flox^ (Asah1^tm1.1Jhkh^) mice [30] were crossed to CD4cre mice and for some experiments additionally to OT-I mice [31] expressing a transgenic T cell receptor recognizing ovalbumin peptide 257-264 (kindly provided by Tetyana Yevsa, Hannover Medical School, Germany). All animal experiments were carried out in accordance with the guidelines of the German Animal Protection Law and were approved by the state authority for nature, environment and customer protection, North Rhine-Westphalia, Germany.

### Cell lines

B16-F1 melanoma cells were cultured in IMDM complete culture medium (IMDM supplemented with 10 % heat-inactivated FCS, 25 µM β-mercaptoethanol and antibiotics (100 U/ml penicillin, 0.1 mg/ml streptomycin)). Cells were maintained in a humidified 5 % CO2 atmosphere at 37°C. Cells were stored in liquid nitrogen and passaged twice before transplantation. Mycoplasma testing was performed every other month by PCR in *in vitro* propagated cultures.

### Tumor transplantation

Tumor cells were harvested and washed twice with PBS. 5 × 10^5^ tumor cells in a volume of 100 µl PBS were injected subcutaneously (s.c.) into the right flank of experimental animals. Tumor volume was calculated using the formula V = (W^2^ x L)/2 [32] based on caliper measurements once tumors have established.

### Amitriptyline treatment

B16-F1 melanoma cells were injected s.c. into 8-12 week old C57BL/6 mice, following mice received 20 mg amitriptyline/kg bodyweight in 100 µl PBS via daily i.p. injection over a period of 13 days.

### Cell Isolation and Activation

Single cell suspensions of splenocytes were generated by rinsing spleens with erythrocyte lysis buffer and washing with PBS supplemented with 2 % FCS and 2 mM EDTA. T cells were isolated from splenocytes either by using the CD4^+^ or CD8^+^ T cell isolation kit (Miltenyi Biotec, Bergisch Gladbach, Germany) according to the manufacturer’s recommendation alone or followed by anti-CD4, anti-CD25, anti-CD8 staining, and cell sorting using an Aria II Cell Sorter (BD Biosciences, Heidelberg, Germany). T cells were stimulated with 1 μg/ml anti-CD3 plate-bound and 1 μg/ml anti-CD28 soluble (both BD Biosciences, Heidelberg, Germany) in IMDM complete culture medium. For exogenous ceramide administration *in vitro* C16 ceramide (Avanti Polar Lipids, Birmingham, USA) solved in 100% EtOH was sonicated for 10 min. The final concentration used for *in vitro* T cell culture was 5 µM.

### Cell isolation from draining lymph nodes (dLN) and tumors

dLN were pestled through a 70-μm cell strainer and washed with PBS containing 2 mM EDTA and 2 % FCS. Tumors were homogenized and pestled through a 70-μm cell strainer and washed with IMDM complete culture medium.

### T cell Differentiation

For Treg differentiation (iTreg) CD4^+^CD25^-^ T cells were stimulated with anti-CD3/anti-CD28 as described above in the presence of 20 ng/ml IL-2 (eBioscience, ThermoFisher Scientific, Langenselbold, Germany) and 5 ng/ml TGF-β1 (R&D Systems, Bio-Techne, Wiesbaden, Germany) for 72 hours.

### Killing Assay

For the generation of antigen-specific cytotoxic lymphocytes (CTLs) splenic CD8^+^ T cells from Asm^flox/flox^/CD4cre/OT-I, Ac^flox/flox^/CD4cre/OT-I mice or the respective littermate controls were cultivated in the presence of irradiated splenocytes, 1 µg/ml OVA-peptide 257-264, 10 ng/ml IL-2, and 20 ng/ml IL-12 for 6 days. As control, cells were stimulated without OVA-peptide 257-264 (non-CTLs). At day 3, cells were split and fresh IMDM complete culture medium supplemented with 10 ng/ml IL-2 was added. CTLs and non-CTLs were isolated from the culture and incubated with OVA-peptide 257-264 loaded CFSE^high^-labeled (2.5 µM CFSE) target and unloaded CFSE^low^-labeled (0.25 µM CFSE) control cells (both splenocytes from WT mice) for 2 or 4 h. Frequencies of target and control populations were analyzed using flow cytometry. Specific killing was calculated as described before [33] using the formula: specific killing [%] = 100-([(CTL^target^/CTL^control^)/(non-CTL^target^/non-CTL^control^)] x 100).

### Antibodies and Flow Cytometry

Anti-CD4, anti-CD8, anti-IFN-γ, anti-CD25, anti-CD44 (BD Biosciences, Heidelberg Germany), anti-CD8, anti-Foxp3, anti-TNF-α, anti-CD69, (eBioscience, ThermoFisher Scientific, Langenselbold, Germany), anti-p-PLCγ, anti-p-ZAP70 (Cell Signaling, Frankfurt am Main, Germany), anti-Granzyme B (Invitrogen, ThermoFisher Scientific, Langenselbold, Germany), and anti-ZAP70 (BioLegend, San Diego, USA) were used as fluorescein isothiocyanate (FITC), pacific blue (PB), phycoerythrin (PE), allophycocyanin (APC), AlexaFluor488 (AF488), AlexaFluor647 (AF647), PE-cyanin 7 (PE-Cy7), or peridinin-chlorophyll protein (PerCp) conjugates. Dead cells were identified by staining with the fixable viability dye eFluor 780 (eBioscience, ThermoFisher Scientific, Langenselbold, Germany). Intracellular staining for Foxp3 and Granzyme B was performed with the Foxp3 staining kit (eBiocience, ThermoFisher Scientific, Langenselbold, Germany) according to the manufacturer’s protocol. IFN-γ and TNF-α expression was measured by stimulating cells with 10 ng/ml phorbol 12-myristate 13-acetate (PMA) and 100 μg/ml ionomycin (both Sigma-Aldrich, München, Germany) for 4 hours in the presence of 5 μg/ml Brefeldin A (Sigma-Aldrich, München. Germany), followed by treatment with 2 % paraformaldehyde and 0.1 % IGEPAL^®^CA-630 (Sigma-Aldrich, München, Germany), and staining with the respective antibody for 30 minutes at 4 °C. Flow cytometric analyses were performed with a LSR II and Canto II instrument using DIVA software (BD Biosciences, Heidelberg Germany).

### Ceramide Quantification by HPLC-MS/MS

Cell suspensions were subjected to lipid extraction using 1.5 ml methanol/chloroform (2:1, v:v) as described [34]. The extraction solvent contained C17 ceramide (C17 Cer) (Avanti Polar Lipids, Alabaster, USA) as internal standard. Chromatographic separations were achieved on a 1290 Infinity II HPLC (Agilent Technologies, Waldbronn, Germany) equipped with a Poroshell 120 EC-C8 column (3.0 × 150 mm, 2.7 µm; Agilent Technologies). MS/MS analyses were carried out using a 6495 triple-quadrupole mass spectrometer (Agilent Technologies) operating in the positive electrospray ionization mode (ESI+) [35]. The following mass transitions were recorded (qualifier product ions in parentheses): *m/z* 520.5 → 264.3 (282.3) for C16 Cer, *m/z* 534.5 → 264.3 (282.3) for C17 Cer, *m/z* 548.5 → 264.3 (282.3) for C18 Cer, *m/z* 576.6 → 264.3 (282.3) for C20 Cer, *m/z* 604.6 → 264.3 (282.3) for C22 Cer, *m/z* 630.6 → 264.3 (282.3) for C24:1 Cer, and *m/z* 632.6 → 264.3 (282.3) for C24 Cer. Peak areas of Cer subspecies, as determined with MassHunter software (Agilent Technologies), were normalized to those of the internal standard (C17 Cer) followed by external calibration in the range of 1 fmol to 50 pmol on column. Determined ceramide amounts were normalized to cell count.

### T cell receptor signaling

For analyzing phosphorylation of TCR signaling molecules by flow cytometry, 5 × 10^5^ splenocytes were left unstimulated or stimulated with 1 μg/ml anti-CD3 and 1 μg/ml anti-CD28 for 5 or 10 minutes, treated with Cytofix/Cytoperm (BD Biosciences, Heidelberg Germany) for 1 hour, and stained with the respective antibodies for 30 minutes at 4°C. For western blot analysis isolated T cells were stimulated for 5 min, washed with PBS, collected in lysis buffer and incubated on ice for 20 min. Afterwards, cells were centrifuged for 5 min at 1200 rpm, the supernatant was collected and protein concentrations were determined as described by Lowry et al. [36]. 30 µg of total protein were diluted in sodium dodecyl sulfate (SDS)-buffer, denatured at 95 °C for 5 min and subjected to sodium dodecyl sulfate polyacrylamide gel electrophoresis (SDS-PAGE). Separated proteins were transferred to a PVDF membrane using the Trans-Blot Turbo RTA Transfer Kit (Bio-Rad Laboratories, Feldkirchen, Germany) according to the manufacturer’s recommendations. The PVDF membrane was blocked with 5 % BSA in TBS-T for 1 h at room temperature. Primary antibodies against p-ZAP70 (Cell Signaling, Frankfurt am Main, Germany, 1:1000) and β-actin (Sigma-Aldrich, Saint Louis, USA, 1:2000) were incubated over night at 4 °C. Secondary anti-rabbit IgG antibody (Sigma-Aldrich, Saint Louis, USA, 1:10,000) was incubated for 1 h at room temperature. Blots were developed using SuperSignal West Femto Maximum Sensitivity Substrate (Thermo Scientific) and signals were detected with a Fusion FX System (Vilber, Eberhardzell, Germany).

### Microscopy

Isolated CD8^+^ T cells were plated in ibidi µ-slides (Ibidi, Gräfelfing, Germany) and stimulated with CD3/CD28 MACSiBead Particles (Miltenyi Biotec, Bergisch Gladbach, Germany) in a 1:1 ratio for 2 h at room temperature. Afterwards, cells were fixed with 4% PFA, blocked with 1 % BSA in PBS for 30 min and stained with anti-ceramide antibody (LSBio, Seattle, USA, 1:20 in 1 % BSA in PBS) for 1 h. Secondary anti-mouse IgM antibody (BioLegend, San Diego, USA, PE-conjugated, 1:1000) and anti-CD3 (BioLegend, San Diego, USA, FITC-conjugated) or anti-TCR beta (Invitrogen, Carlsbad, USA, FITC-conjugated, 1:50) antibodies were diluted in 1 % BSA in PBS and incubated for 30 min at room temperature. Stained cells were mounted using Fluorescence Mounting Medium (Dako, California, USA) and visualized with a Biorevo BZ-9000 fluorescence microscope (Keyence, Itasca Illinois, USA).

### CD4^+^ T cell depletion

For depletion of CD4^+^ T cells during tumorigenesis, 200 μg anti-mouse CD4-depleting antibody (clone GK1.5; BioXcell, Lebanon, USA) was injected intraperitoneal on days -1, 3, 6, 9, and 12 after tumor cell transplantation.

### RNA isolation, cDNA synthesis and qRT-PCR

RNA was isolated using the NucleoSpin RNA XS Kit (Macherey-Nagel, Düren, Germany) according to the manufacturer’s instructions. 100 ng of RNA was reversed transcribed using M-MLV Reverse Transcriptase (Promega, Mannheim, Germany) with dNTPs (Bio-Budget, Krefeld, Germany), Oligo-dT mixed with Random Hexamer primers (both Invitrogen, Frederick Maryland, USA). Quantitative real-time PCR was performed using the Fast SYBR Green Master Mix (Thermo Fisher Scientific, Braunschweig, Germany) and a 7500 Fast Real-Time PCR System (Thermo Fisher Scientific, Darmstadt, Germany). Samples were measured as technical duplicates. Expression levels were normalized against ribosomal protein S9 (RPS9). Following primer sequences were used: Asm (Smpd1) CTG TCA GCC GTG TCC TCT TCC TTA, GGG CCC AGT CCT TTC AAC AG, Ac (Asah1) TTC TCA CCT GGG TCC TAG CC, TAT GGT GTG CCA CGG AAC TG, RPS9 CTG GAC GAG GGC AAG ATG AAG C, TGA CGT TGG CGG ATG AGC ACA.

### Statistical Analysis

Statistical analyses were calculated using Graph Pad Prism Software (Graph Pad Software, La Jolla, CA). To test for Gaussian distribution, D’Agostino-Pearson omnibus and Shapiro-Wilk normality tests were used. If data passed normality testing, paired or unpaired Student’s t-test was performed, otherwise Mann-Whitney U-test was used for unpaired data. Differences between two or more groups with different factors were calculated using 2way ANOVA followed by Sidak’s post-test. Statistical significance was set at the levels of *p<0.05, **p<0.01, ***p<0.001, and ****p<0.0001.

## Results

### Asm-deficient mice show enhanced tumor growth accompanied by reduced T cell activation

To analyze the impact of Asm activity on T cell function during tumorigenesis, we transplanted B16-F1 melanoma cells into Asm-deficient mice or WT littermates. We observed significantly accelerated tumor growth rates in Asm-deficient mice compared to WT mice as previously described [16] (Fig. 1A). Subsequently, we analyzed T cell frequencies in tumor draining lymph nodes (dLN) and tumors as well as the activation status of tumor infiltrating lymphocytes (TILs). We detected lower frequencies of CD4^+^ and CD8^+^ T cells in dLNs but increased frequencies of Tregs in tumor bearing Asm-deficient mice compared to WT mice (Fig. 1B). Furthermore, CD4^+^ and CD8^+^ TILs showed a reduced expression of IFN-γ and CD44 (Fig. 1C), indicating a decreased T cell response in Asm-deficient mice during tumorigenesis. These results suggest that elevated tumor growth in Asm-deficient mice correlates with an insufficient T cell response.

**Fig. 1.**
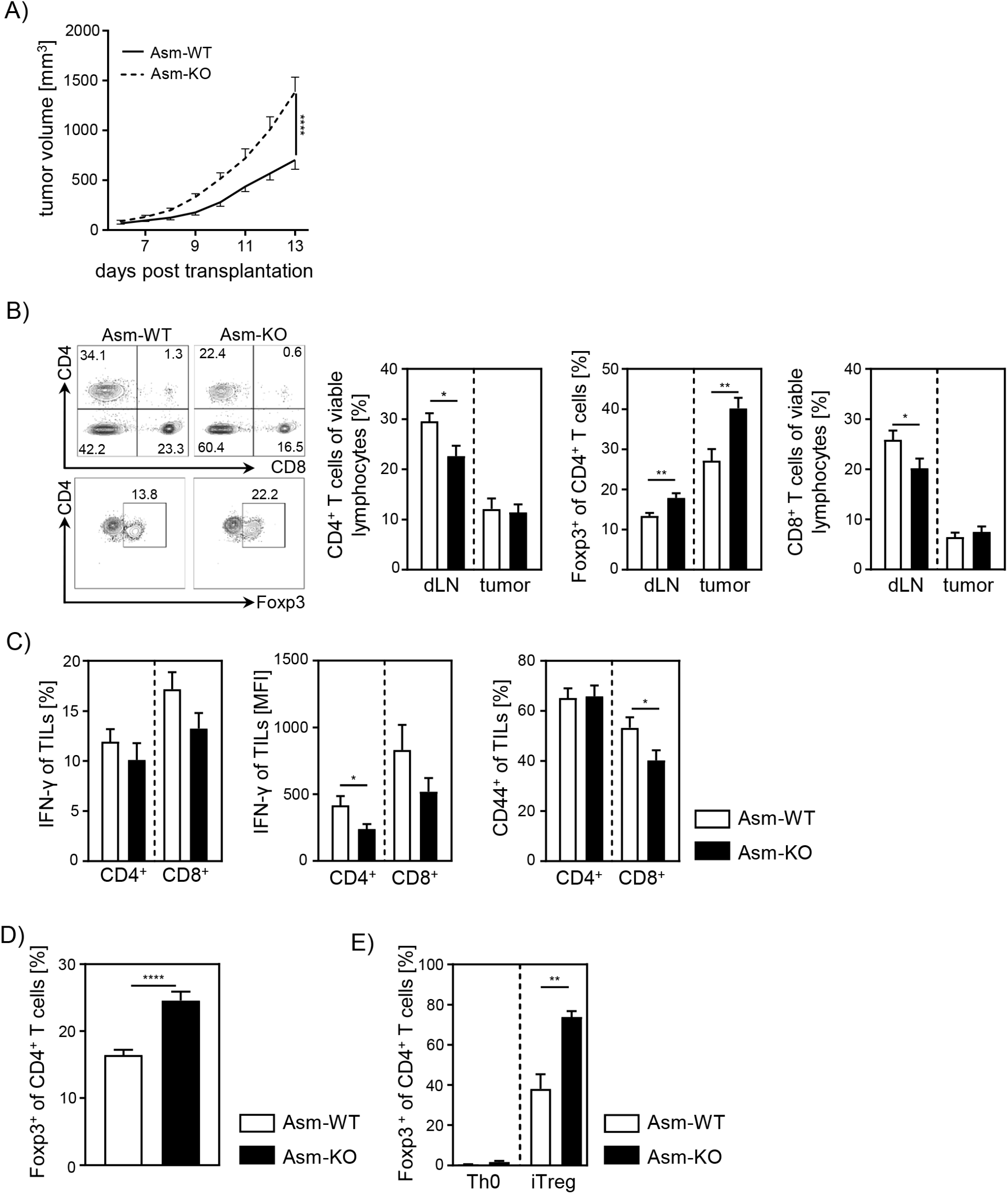
Ablation of Asm results in decreased T cell activation and enhanced tumor growth. (A) B16-F1 melanoma cells were transplanted into Asm-KO mice and WT littermates. Tumor volume was monitored daily once tumors were palpable (n= 12-18). (B) Frequencies of CD4^+^ T cells, CD4^+^Foxp3^+^ Tregs, and CD8^+^ T cells within draining lymph nodes (dLN) and tumor of Asm-KO and Asm-WT mice were determined by flow cytometry. Representative contour plots (dLN) are shown in the left panel. (C) IFN-γ, and CD44 expression of CD4^+^ and CD8^+^ tumor-infiltrating lymphocytes (TILs) in tumor-bearing mice. (D) Percentages of Foxp3^+^ Tregs of CD4^+^ T cells within spleen of naïve Asm-KO and Asm-WT mice were determined by flow cytometry (n= 16-17). (E) Sorted CD4^+^CD25^-^ T cells from Asm-KO and Asm-WT mice were stimulated with anti-CD3 and anti-CD28 in the presence of IL-2 and TGF-β1 (iTreg). Respective controls (Th0) were only stimulated with anti-CD3 and anti-CD28 antibodies. After 3 days, Treg differentiation was analyzed by Foxp3 expression (n= 3-4). Results from 2 to 4 independent experiments are depicted as mean ± SEM. Statistical analysis was performed by 2way ANOVA with Sidak’s multiple comparisons or Student’s t-test. (*p < 0.05, **p < 0.01, ****p < 0.0001).

Moreover, we investigated whether pharmacological inhibition of Asm also affects the T cell response during tumorigenesis in a similar way. Therefore, we treated tumor-bearing C57BL/6 mice with amitriptyline by daily i.p. injection. Indeed, we detected enhanced tumor progression in amitriptyline-treated mice compared to tumor-bearing mice that received the vehicle. This was accompanied by reduced activation of TILs in terms of IFN-γ and CD44 expression in amitriptyline-treated C57BL/6 mice compared to PBS-treated mice (Supplemental Figure S1A, B).

Next, we analyzed the effect of Asm deficiency on the Foxp3^+^ Treg subpopulation *in vitro*, since we observed elevated frequencies of Tregs in dLN and tumors of tumor-bearing Asm-deficient mice. In accordance with results from Hollmann et. al. [26] and our own study of Asm-overexpressing T cells [27], we detected elevated Treg frequencies in spleens of naïve Asm-KO mice as compared to controls (Fig. 1D). Moreover, induction of Tregs *in vitro* revealed an improved capacity of CD4^+^CD25^-^ T cells to differentiate into Tregs when Asm activity is absent (Fig. 1E). In summary, these data provide evidence that ceramide generation by Asm activity in CD4^+^ T cells interferes with Treg differentiation.

### CD4^+^ T cell depletion in tumor-bearing Asm-deficient mice reveals CD8^+^ T cell dysfunction

Next, we asked whether the observed increase in relative numbers of Tregs contribute to the insufficient CD8^+^ T cell response accompanied by enhanced tumor growth in Asm-deficient mice. For this purpose, we depleted CD4^+^ T cells from Asm-deficient and Asm-WT mice and transplanted B16-F1 melanoma cells. Interestingly, depletion of CD4^+^ T cells reduced tumor growth in both Asm-WT and Asm-KO mice. Still, tumors of CD4^+^ T cell-depleted Asm-deficient mice showed higher tumor growth rates than CD4^+^ T cell-depleted Asm-WT mice (Fig. 2A). Remarkably, depletion of CD4^+^ T cells in Asm-WT mice abolished tumor progression almost completely. Using flow cytometry analysis, we confirmed the successful depletion of CD4^+^ T cells in dLN and tumor tissue and observed an increase of CD8^+^ T cell frequencies and numbers, which was significantly less pronounced in dLNs and tumors of Asm-KO mice (Fig. 2B). Strikingly, analysis of CD8^+^ TILs from Asm-deficient mice revealed reduced expression of activation-associated molecules like IFN-γ, CD44 and granzyme B as compared to Asm-WT mice after CD4^+^ T cell depletion (Fig. 2C). These results indicate that Asm-deficiency interferes with CD8^+^ T cell responses in tumor-bearing mice independent of CD4^+^ T cells.

**Fig. 2.**
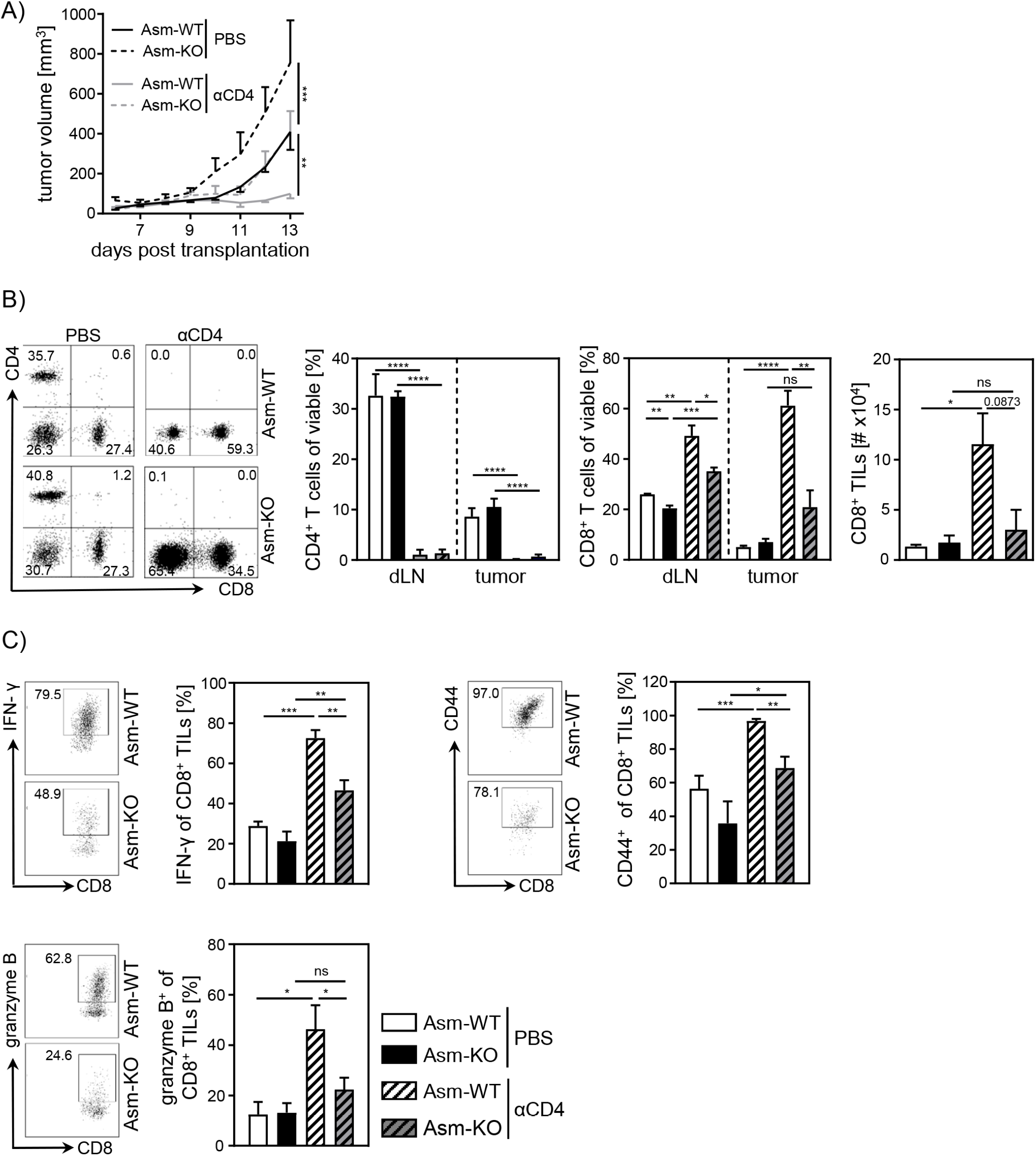
Impaired CD8^+^ T cell function in Asm-deficient tumor-bearing mice upon CD4^+^ T cell depletion. CD4^+^ T cells were depleted from Asm-WT and Asm-KO mice by repeated i.p. injection of anti-CD4 depleting antibody. Control groups received PBS. B16-F1 tumor cells were transplanted one day later s.c. (A) Tumor volume was determined after tumor establishment based on caliper measurements (n= 3-7). (B) Frequencies of CD4^+^ and CD8^+^ T cells in dLN and tumor and (C) expression of IFN-γ, CD44, and granzyme B of CD8^+^ tumor-infiltrating lymphocytes (TILs) were analyzed using flow cytometry. Representative dots plots are shown in the left panel. Results from 2 independent experiments are depicted as mean ± SEM. Statistical analysis was performed by 2way ANOVA with Sidak’s multiple comparisons or Student’s t-test. (*p < 0.05, **p < 0.01, ***p < 0.001, ****p < 0.0001).

### T cell-specific ablation of Asm reduces CD8^+^ T cell activation *in vitro*

To gain further insights into the role of cell-intrinsic Asm activity in CD8^+^ T cells, we analyzed the phenotype and function of Asm-deficient CD8^+^ T cells *in vitro*. To exclude an impact of other Asm-deficient cells present in Asm-KO mice, we made use of Asm^flox/flox^/CD4cre (Asm^flox/flox^/CD4cre^tg^) mice, which lack Asm expression specifically in T cells (Fig. 3A, Supplement 2A). This results in significantly reduced ceramide levels in unstimulated, as well as anti-CD3/anti-CD28 stimulated CD8^+^ and CD4^+^ T cells compared to T cells from control littermates (Asm^flox/flox^/CD4cre^wt^) (Fig. 3B, Supplemental Figure S2B). Consistent with decreased T cell activation observed in tumor-bearing Asm-KO mice, *in vitro* stimulation of sorted CD8^+^ T cells from Asm^flox/flox^/CD4cre mice led to reduced expression of early T cell activation-associated molecules. This was reflected by lower CD25, CD69 and CD44 expression among Asm-deficient CD8^+^ T cells compared to Asm-proficient CD8^+^ T cells after 24 hours of stimulation (Fig. 3C). Moreover, co-cultivation of antigen-specific CTLs, generated from Asm^flox/flox^/CD4cre/OT-I mice, together with OVA-loaded target cells revealed a reduced killing capacity of Asm-deficient CD8^+^ T cells (Fig. 3D). Well in line, CD8^+^ T cells from Asm^flox/flox^/CD4cre mice showed reduced granzyme B expression in response to TCR stimulation (Fig.3E). Strikingly, this phenotype was partially rescued by the addition of exogenous C16 ceramide during stimulation (Fig. 3F). Altogether, these data indicate that ceramide is important for effective CD8^+^ T cell responses, at least *in vitro*. In addition to the effect of ceramide on CD8^+^ T cell function, we detected a reduced capacity of CD4^+^ T cells from Asm^flox/flox^/CD4cre mice to differentiate into Th1 cells, but to a higher extent into Tregs *in vitro* (Supplemental Figure S2C, D), which is in line with our previous results from Asm-KO mice (Fig. 1E).

**Fig. 3.**
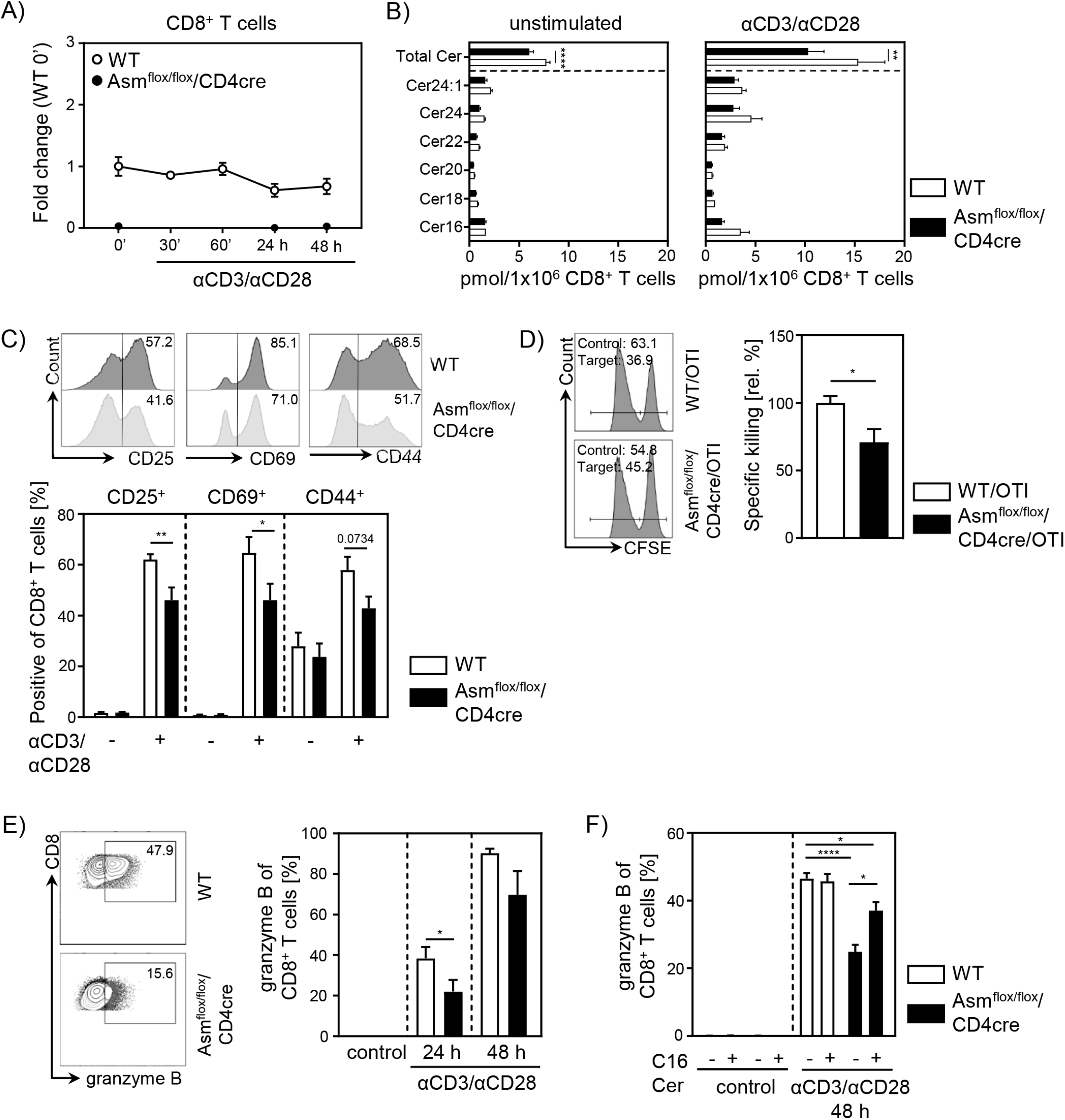
T cell specific Asm deficiency leads to reduced CD8^+^ T cell activation *in vitro*. Isolated CD8^+^ T cells from Asm^flox/flox^/CD4cre mice and WT littermates where either left unstimulated or stimulated with anti-CD3 and anti-CD28 for indicated time points. (A) mRNA expression of Asm (*Smpd1*) following activation was analyzed by RT-qPCR (n=4-8). (B) Ceramide levels of CD8^+^ T cells were determined by mass spectrometry (n= 4). (C) Expression of CD25, CD69, and CD44 was analyzed by flow cytometry (n= 6-7). Representative histograms for are shown in the upper panel. (D) Ova-specific CTLs were generated and incubated with Ova-peptide 257-264 loaded CFSE^high^-labeled target and unloaded CFSE^low^-labeled control cells. Specific killing was evaluated by frequencies of target and control populations determined by flow cytometry (n= 6-7). Representative histograms are shown in the left panel. (E) Frequencies of granzyme B-expressing CD8^+^ T cells from Asm^flox/flox^/CD4cre mice and WT littermates without and (F) in the presence of C16 ceramide were analyzed by flow cytometry (n= 4-8). Representative contour plots are shown in the left panel. Results from 2 to 4 independent experiments are depicted as mean ± SEM. Statistical analysis was performed by 2way ANOVA with Sidak’s multiple comparisons, Mann-Whitney U-test or Student’s t-test. (*p < 0.05, **p < 0.01, ****p < 0.0001).

### Cell-intrinsic Asm activity determines T cell activation during tumorigenesis

To investigate whether T cell-specific Asm ablation has also an impact on T cell responses during tumorigenesis *in vivo*, we transplanted B16-F1 melanoma cells into Asm^flox/flox^/CD4cre mice and observed significantly higher tumor growth rates in Asm^flox/flox^/CD4cre mice compared to control mice (Fig. 4A). Again, we analyzed T cell frequencies and T cell activation in dLNs and TILs. In line with results from Asm-KO mice (Fig. 1B), we detected elevated percentages of tumor infiltrating Foxp3^+^ Tregs as well as reduced frequencies of CD4^+^ and CD8^+^ T cells in dLNs from Asm^flox/flox^/CD4cre mice compared to control littermates (Fig. 4B). Moreover, absolute cell numbers of intratumoral T cells were reduced in Asm^floxflox^/CD4cre mice compared to WT littermates.TILs from T cell-specific Asm-deficient mice showed decreased expression of IFN-γ and TNF-α, as well as granzyme B indicating a reduced anti-tumoral T cell response (Fig. 4C). These results provide evidence that cell-intrinsic Asm activity has an impact on T cell responses *in vitro* and during ongoing immune responses *in vivo*.

**Fig. 4.**
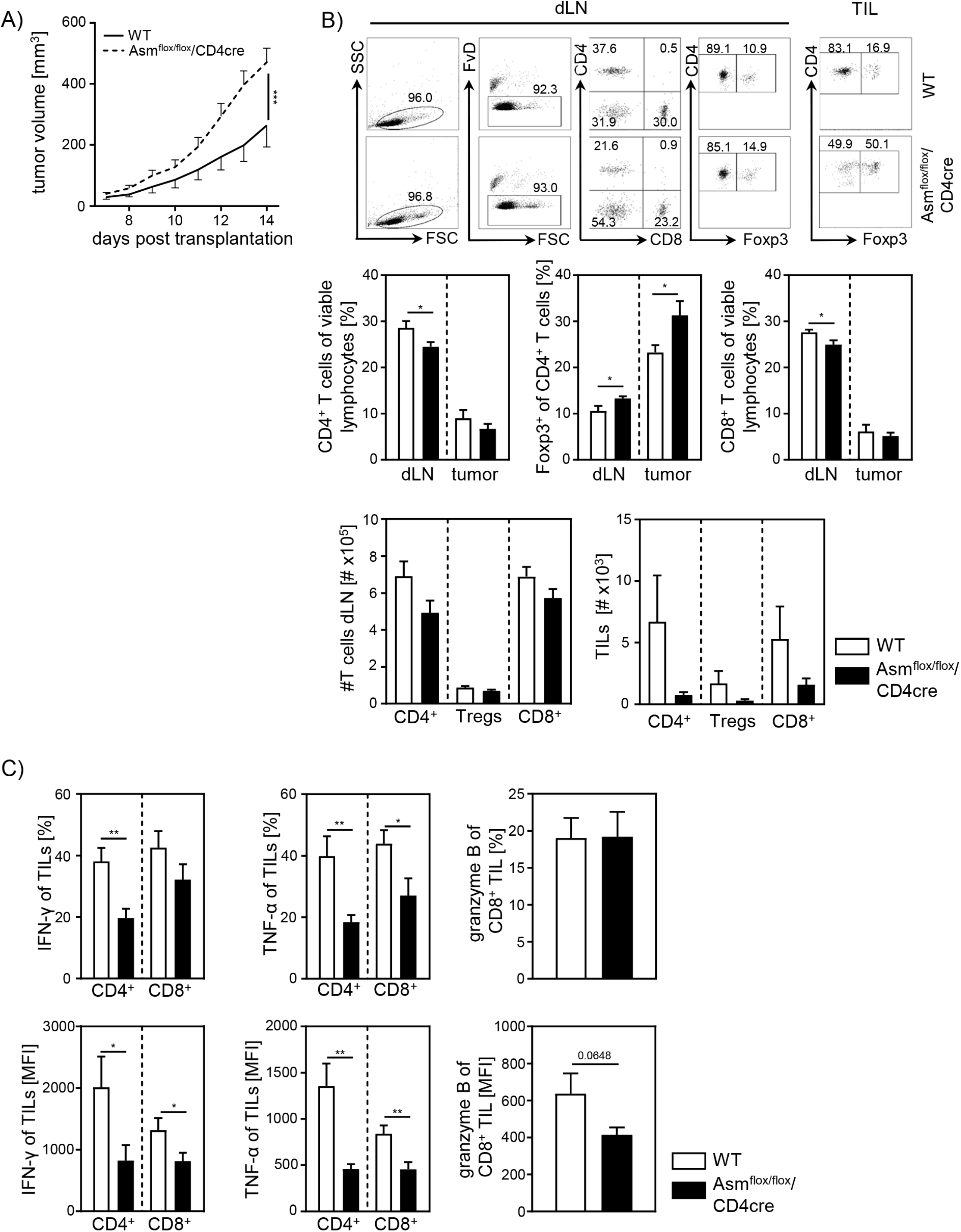
Cell-intrinsic Asm activity determines CD8^+^ T cell activation *in vivo*. (A) B16-F1 melanoma cells were transplanted into Asm^flox/flox^/CD4cre mice and WT littermates and tumor growth was monitored when tumors reached a detectable size (n= 12-16). (B) Percentages of CD4^+^ T cells, Foxp3^+^ Tregs, and CD8^+^ T cells within dLN and tumor were determined by flow cytometry and absolute cell numbers were calculated. Representative dot plots are shown in the upper panel. (C) Expression of IFN-γ, TNF-α, and granzyme B of TILs were determined by flow cytometry. (Results from 4 independent experiments are depicted as mean ± SEM. Statistical analysis was performed by 2way ANOVA with Sidak’s multiple comparisons, Mann-Whitney U-test or Student’s t-test. (*p < 0.05, **p < 0.01, ***p < 0.001).

### Ceramide accumulates at TCR synapsis

The previous experiments indicate that ceramide generation by Asm activity is involved in T cell function. In order to elucidate the subcellular localization of ceramide, we performed fluorescence microscopy of CD8^+^ T cells stimulated with CD3/CD28 MACSiBead Particles and stained for ceramide, CD3, and TCR-beta. Indeed, ceramide accumulates at the contact site between T cell and particle. Strikingly, ceramide co-localizes with CD3 (Fig. 5A) and TCR beta, respectively (Fig. 5B). This co-localization of TCR and ceramide suggests an involvement ceramide in TCR signaling.

**Fig. 5.**
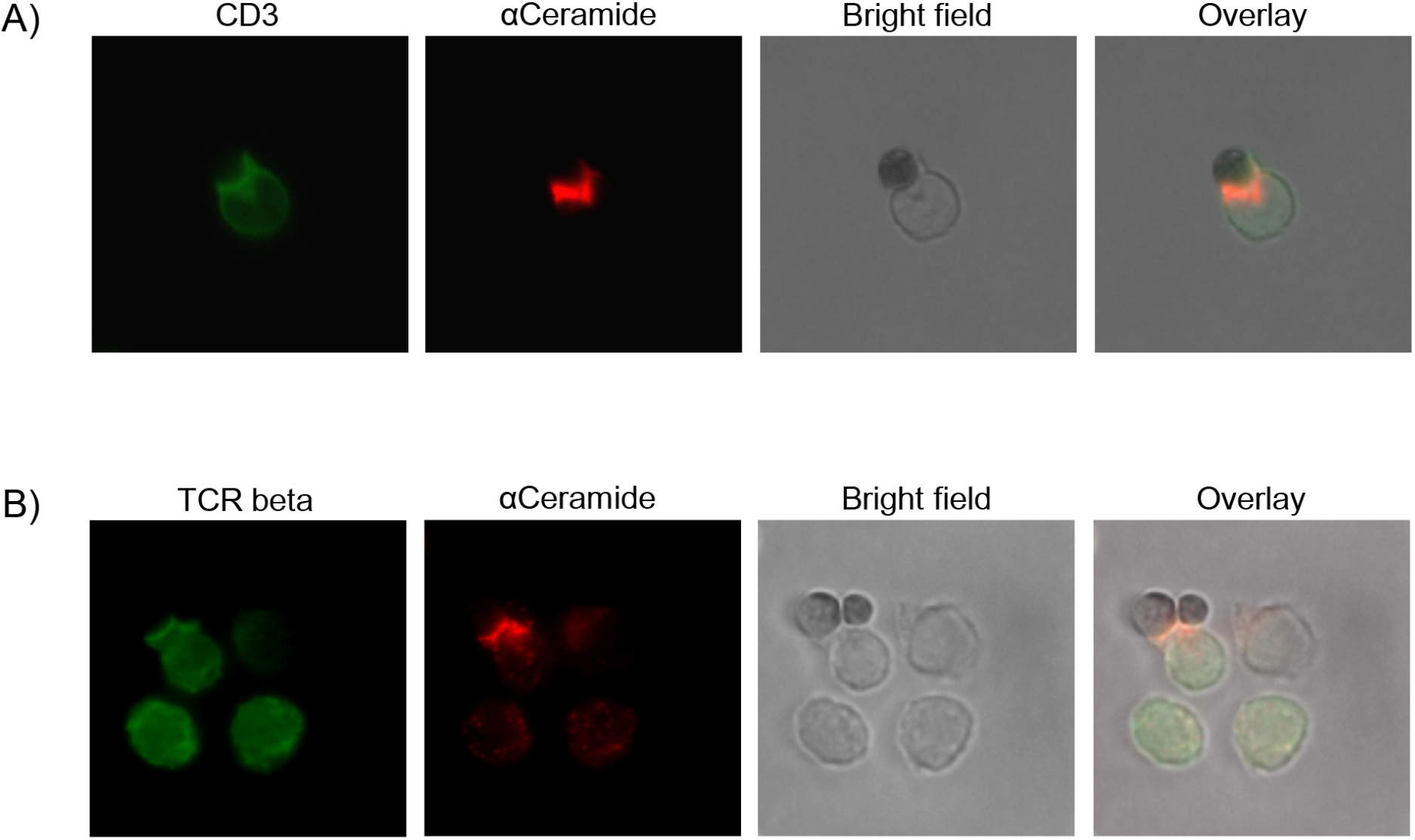
Ceramide co-localizes with CD3 and TCR. CD8^+^ T cells were isolated and stimulated with CD3/CD28 MACSiBead Particles for 2h and stained for ceramide (red) and (A) CD3 or (B) TCR beta (green). Cells were visualized using a Biorevo BZ-9000 fluorescence microscope.

### Elevated ceramide generation in Ac-deficient CD8^+^ T cells results in enhanced cytotoxic function *in vitro*

To further validate the important role of ceramide in CD8^+^ T cell responses we made use of Ac^flox/flox^/CD4cre mice. In these mice, T cells are deficient for Ac, the enzyme that catalyzes the hydrolysis of ceramide into sphingosine. As validated by qPCR both, CD8^+^ and CD4^+^ T cells lack Ac expression (Fig. 6A, Supplemental Figure S3A), resulting in elevated ceramide concentrations in unstimulated and anti-CD3/anti-CD28 T cells from Ac^flox/flox^/CD4cre (Ac^flox/flox^/CD4cre^tg^) mice compared to control littermates (Ac^flox/flox^/CD4cre^wt^) (Fig. 6B, Supplemental Figure S3B). Since we observed a co-localization between ceramide and CD3 and TCR-beta upon stimulation, respectively (Fig 5), we wondered whether elevated ceramide levels in Ac-deficient T cells might correlate with TCR signaling. Therefore, we isolated splenocytes from Ac^flox/flox^/CD4cre and control mice and stimulated them with anti-CD3/anti-CD28 *in vitro*. Indeed, CD8^+^ and CD4^+^ T cells from Ac^flox/flox^/CD4cre mice showed significantly elevated phosphorylation of the TCR signaling molecules ZAP70 and PLCγ compared to control CD8^+^ T cells (Fig. 6C, Supplemental Figure S3C). The elevated phosphorylation of ZAP70 was further confirmed by western blot analysis of isolated Ac-deficient CD8^+^ T cells (Fig. 6D). Well in line with these data, stimulation of CD8^+^ T cells from Ac^flox/flox^/CD4cre mice led to an increased expression of granzyme B compared to CD8^+^ T cells from control littermates (Fig. 6E). Strikingly, CTLs from Ac^flox/flox^/CD4cre/OT-I mice showed an improved killing capacity in comparison to control cells (Fig. 6F). Although Ac ablation had no impact on Th1 differentiation in regard to IFN-γ expression, CD4^+^ T cells from Ac^flox/flox^/CD4cre mice showed an enhanced expression of granzyme B under Th1-polarizing conditions *in vitro* (Supplemental Figure S3D).

**Fig. 6.**
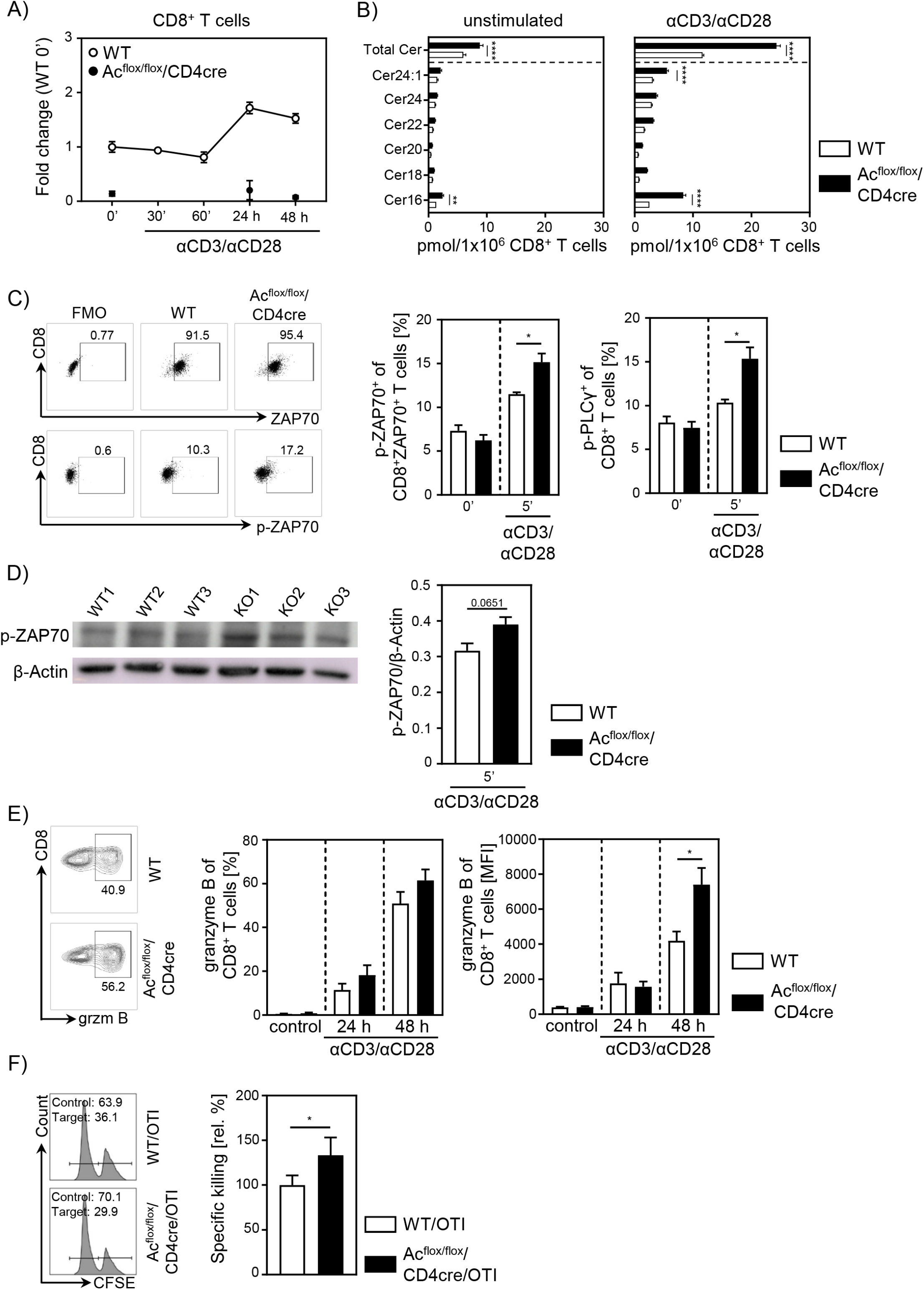
Ac-deficient CD8^+^ T cells have elevated ceramide levels and show increased activation *in vitro*. (A) Isolated CD8^+^ T cells from Ac^flox/flox^/CD4cre mice and WT littermates where either left unstimulated or stimulated with anti-CD3 and anti-CD28 for indicated time points. mRNA expression of Ac (*Asah1*) following activation was analyzed by RT-qPCR (n= 3-4). (B) Ceramide levels of CD8^+^ T cells were determined by mass spectrometry (n= 4). (C) For TCR signaling analysis, splenocytes from Ac^flox/flox^/CD4cre and WT mice were left unstimulated (0’) or stimulated with anti-CD3 and anti-CD28 for 5 (5’) min. Afterwards, samples were analyzed for phospho-ZAP70 of gated ZAP70^+^CD8^+^ T cells and phosphor-PLCγ of gated CD8^+^ T cells by flow cytometry (n= 4). Representative dot plots and FMOs for ZAP70 are shown in the left panel. (D) Western blot analysis of phospho-ZAP70 expression of CD8^+^ T cells from Ac^flox/flox^/CD4cre and WT mice after 5 min of stimulation with anti-CD3 and anti-CD28 (n= 3). (E) CD8^+^ T cells were left untreated as control or stimulated for 24 or 48 h and analyzed for granzyme B expression by flow cytometry (n= 5-8). Representative contour plots are shown in the left panel. (F) Specific killing of antigen-specific CTLs from Ac^flox/flox^/CD4cre/OTI mice and WT controls was assessed (n= 8-9). Representative histograms are shown in the left panel. Data are depicted as mean ± SEM. Statistical analysis was performed by 2way ANOVA with Sidak’s multiple comparisons, Mann-Whitney U-test or Student’s t-test. (*p < 0.05, **p < 0.01, ****p < 0.0001).

### T cell-specific Ac ablation promotes anti-tumoral immune response

Since our data indicate an important role for cell-intrinsic ceramide in T cell activation *in vitro*, we next analyzed the effect of accumulating ceramide in T cells *in vivo*. Thus, we transplanted B16-F1 melanoma cells into Ac^flox/flox^/CD4cre mice and control littermates. In contrast to Asm^flox/flox^/CD4cre mice, Ac^flox/flox^/CD4cre mice showed a significant reduction in tumor size compared to control mice (Fig. 7A). Although mice did not differ regarding T cell frequencies or numbers (Fig. 7B), we detected an elevated T cell activation in Ac^flox/flox^/CD4cre tumor-bearing mice. This was reflected by increased IFN-γ and granzyme B expression of CD4^+^ and CD8^+^ TILs in comparison to control mice (Fig. 7C). From these results, we conclude that enhancing cell-intrinsic ceramide by ablation of Ac activity promotes the T cell function, whereas reduced ceramide levels due to loss of Asm activity is accompanied by a decrease in the activity of T cells.

**Fig. 7.**
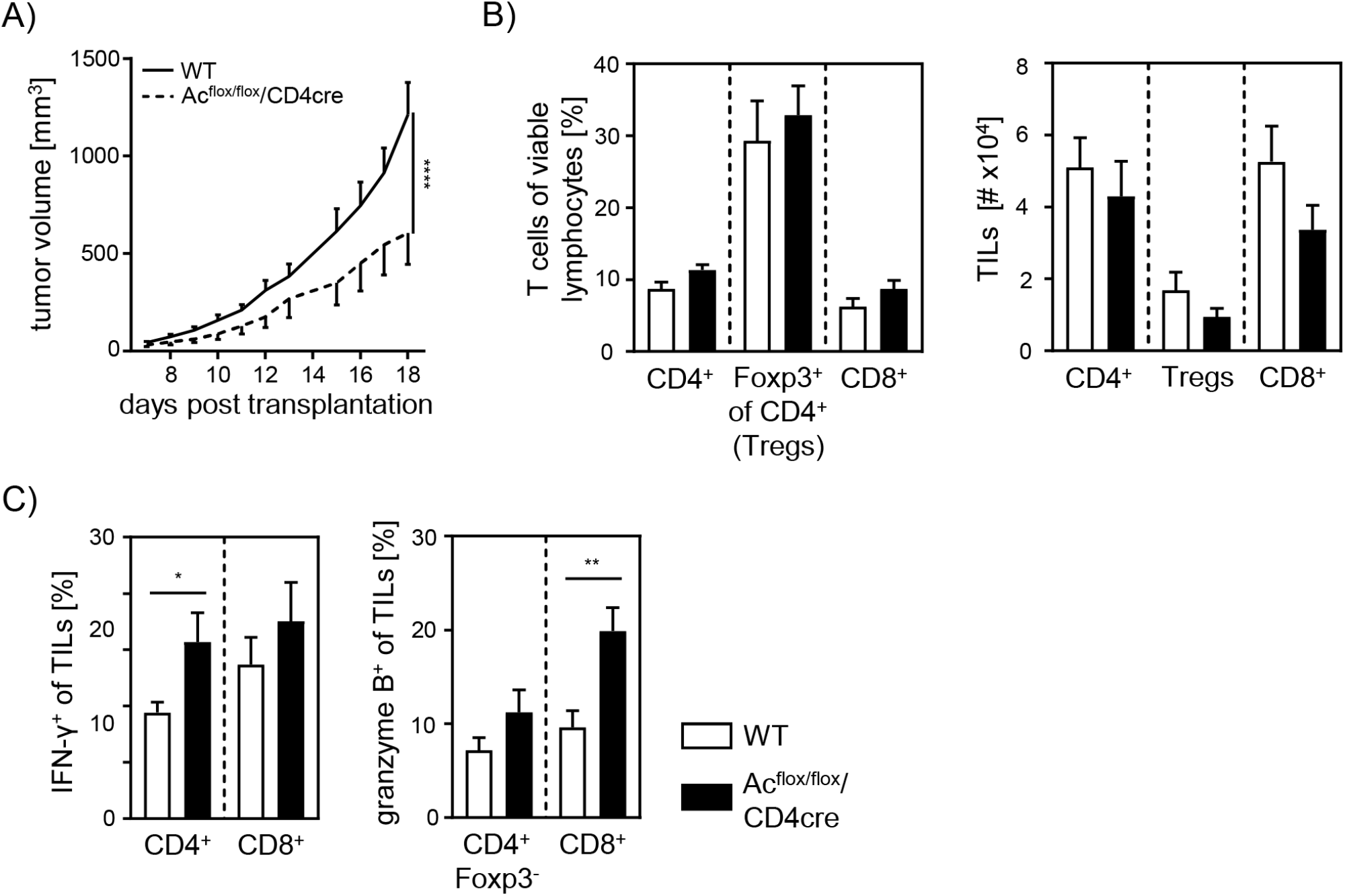
Elevated anti-tumor immune response in T cell-specific Ac-deficient mice. (A) B16-F1 melanoma cells were transplanted into Ac^flox/flox^/CD4cre mice and WT littermates. Tumor volume was monitored once tumors have been established (n= 8-10). (B) Percentages of CD4^+^ T cells, Foxp3^+^ Tregs, and CD8^+^ T cells were determined by flow cytometry and absolute cell numbers were calculated. (C) Frequencies of IFN-γ^+^ and granzyme B^+^ TILs were determined by flow cytometry. Results from 2 independent experiments are depicted as mean ± SEM. Statistical analysis was performed by 2way ANOVA with Sidak’s multiple comparisons or Student’s t-test. (*p < 0.05, **p < 0.01, ****p < 0.0001).

## Discussion

In the present study, we investigated the impact of cell-intrinsic Asm and Ac activity on the phenotype and function of CD4^+^ as well as CD8^+^ T cells *in vitro*, and during tumorigenesis *in vivo*. We identified a correlation between ceramide levels and T cell activity. Reduced ceramide content due to loss of Asm activity triggered Treg induction and interfered with effector T cell responses, whereas elevated ceramide concentrations induced by ablation of Ac expression resulted in enhanced T cell activation. Strikingly, cell-intrinsic ceramide levels also correlated with anti-tumoral immune responses and tumor growth in T cell-specific Asm-deficient or Ac-deficient mice.

Several studies already described Asm as regulator of CD4^+^ T cell function and differentiation [25], [26], [28]. Well in line, we demonstrated that cell-intrinsic Asm activity determines T cell activation and differentiation. However, the mechanism that triggers Asm activity in T cells is still controversial. Mueller et. al. postulated that CD28 stimulation induces Asm activity in T cells, whereas co-stimulation of CD3 and CD28 does not [37]. In contrast, Asm activation has been described in isolated human CD4^+^ T cells in response to CD3/CD28 co-stimulation [28], while Wiese et al. identified CD28 co-stimulation to be required for enhanced human CD4^+^Foxp3^+^ Treg frequencies upon Asm inhibition [38]. In accordance, we detected significantly elevated ceramide concentrations upon anti-CD3/anti-CD28 treatment of WT CD4^+^ T cells, which were substantially reduced in Asm-deficient T cells, suggesting that co-stimulation of CD3/CD28 is important for the induction of Asm activity in CD4^+^ T cells.

Others and we demonstrated that transplantation of melanoma cells into Asm-deficient mice results in accelerated tumor growth. As one underlying mechanism, apoptosis-resistant endothelial cells has been proposed [39]. Here, we provide evidence that Asm-deficiency in T cells contribute to an impaired anti-tumoral immune response resulting in loss of tumor growth control. We detected elevated percentages of Foxp3^+^ Tregs accompanied with reduced expression of activation-associated molecules of effector T cells from tumor-bearing Asm-deficient mice, amitriptyline-treated mice and importantly, in T cell-specific Asm-deficient mice, in contrast to the respective controls. These results indicate that the T cell-intrinsic Asm activity regulates T cell function and differentiation most likely via the generation of ceramide.

It is well established that Tregs infiltrate the tumor tissue and interfere with an effective local immune response [22], [23], [40], [41]. Moreover, a higher ratio of Tregs to CD8^+^ effector T cells correlates with poorer disease outcome in cancer [42]–[44]. To dissect whether enhanced Treg frequencies from Asm-deficient mice are responsible for the enhanced tumor growth, we depleted CD4^+^ T cells from tumor-bearing mice. Interestingly, CD4^+^ T cell depletion resulted in reduced tumor growth in both AsmWT and Asm-KO mice. This is in line with studies by Ueha et. al., who observed a reduction of tumor growth after CD4^+^ T cell depletion [45]. However, the tumor growth in Asm-deficient mice was still significantly enhanced compared to WT littermates after CD4^+^ T cell depletion. Subsequent analysis of CD8^+^ T cells revealed reduced activation of CD8^+^ T cells from tumor-bearing Asm-deficient mice. These results provide evidence, that Asm activity has an impact not only on CD4^+^ T cell subsets, but also on CD8^+^ T cell function during tumorigenesis. A reduced capacity to release cytotoxic granules by antigen-specific CD8^+^ T cells from LCMV-infected Asm-deficient mice has already been described by Herz et. al. [24]. In accordance, we also observed decreased frequencies of granzyme B producing CD8^+^ TILs in tumor-bearing Asm-deficient mice, after CD4^+^ T cell depletion. Further analysis of CD8^+^ T cells from T cell-specific Asm-deficient mice (Asm^flox/flox^/CD4cre) revealed a reduced activation upon stimulation *in vitro* compared to WT controls. Strikingly, Asm-deficiency was associated with a lowered *in vitro* killing capacity of CD8^+^ CTLs, suggesting that the ceramide content could regulate cytotoxic CD8^+^ T cell function. Indeed, Asm-deficient CD8^+^ T cells with reduced ceramide concentrations showed an impaired granzyme B production. Interestingly, this phenotype could partially be rescued by the addition of C16 ceramide *in vitro*. Validating the impact of ceramide on CD8^+^ T cell responses, we used Ac^flox/flox^/CD4cre mice with T cell-specific ablation of Ac expression resulting in increased ceramide concentrations in CD8^+^ T cells. Well in line with results from Asm-deficient CD8^+^ T cells, we demonstrated, at least to our knowledge, for the first time, that Ac-deficient CD8^+^ T cells show increased granzyme B expression upon stimulation and importantly, elevated *in vitro* killing activity. This phenotype correlated with a facilitated anti-tumoral T cell response leading to reduced tumor growth rates of Ac^flox/flox^/CD4cre mice transplanted with B16-F1 melanoma cells in contrast to WT littermates. These results indicate that ceramide concentrations may determine T cell activity and function *in vitro* and *in vivo*. In previous studies intracellular sphingosine-1-phosphate (S-1-P) has been shown to reduce anti-tumor functions of T cells [46], [47]. To exclude that the modulated anti-tumoral T cell response in Asm^flox/flox^/CD4cre and Ac^flox/flox^/CD4cre mice, respectively is not caused by altered S-1-P concentrations, we quantified its abundance in CD3/CD28 stimulated T cells. However, S-1-P concentrations were under the detection limit, emphasizing, that indeed ceramide modulates the anti-tumoral T cell response more likely that S-1-P levels.

Our results demonstrate that ablation of Ac in CD8^+^ T cells from Ac^flox/flox^/CD4cre mice led to a facilitated phosphorylation of ZAP-70 and PLCγ suggesting that ceramide, most likely as ceramide-enriched platforms in the plasma membrane, modulates the TCR signaling pathway. Indeed, by using fluorescence microscopy, we were able to reveal a co-localization of ceramide and TCR in stimulated CD8^+^ T cells, supporting our hypothesis that the ceramide level is involved in the strength of TCR-induced signaling pathway. In accordance Bai and colleagues observed reduced phosphorylation of different TCR signaling molecules in stimulated human CD4^+^ T cells in the presence of the Asm inhibitor imipramine [28], further suggesting that TCR signaling is affected by ceramide. In addition, one might speculate about a positive feedback regulation between ceramide and T cell activation since TCR stimulation of CD4^+^ as well as CD8^+^ T cells resulted in elevated ceramide concentration, which in turn seems to strengthen the TCR signaling cascade.

Overall, our study provides evidence that the cell-intrinsic ceramide content is regulated by the activity of Asm and Ac and associated with CD4^+^ as well as CD8^+^ T cell function *in vitro* and *in vivo*. Thereby, the sphingolipid metabolism represents a potential therapeutic target for improving anti-tumoral T cell responses during tumorigenesis. However, the use of drugs modulating ceramide generating enzymes like amitriptyline or other FIASMAs should be carefully reflected in cancer patients. Nevertheless, genetic engineering of T cells by manipulating the expression of sphingolipid metabolizing enzymes and using them for cancer therapy could be a potential approach for cancer therapy.

## Acknowledgement

We kindly thank Sina Luppus for excellent technical assistance and Witold Bartosik and Christian Fehring for cell sorting. Moreover, we thank Daniel Herrmann for the help with the HPLC-MS analyses of ceramides. This work was supported by the Deutsche Forschungsgemeinschaft (DFG -GRK2098 to AMW, JB, KAB, EG and WH, and GRK1949 to AMW, JB, and WH).

---

**Supplemental Fig. S1.**
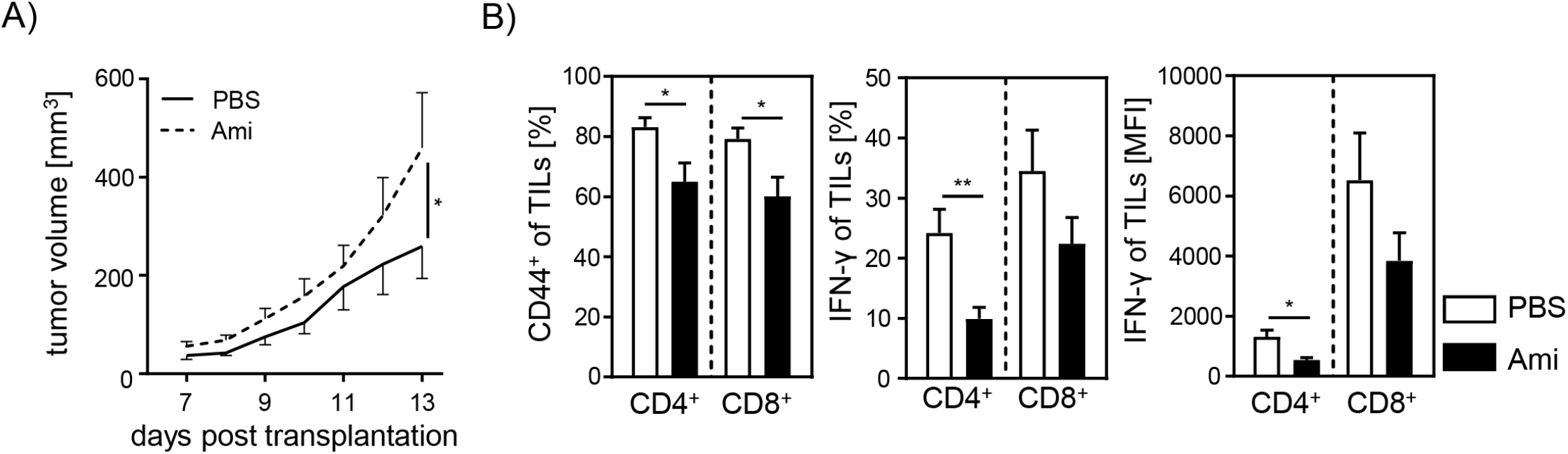
(A) B16-F1 melanoma cells were injected into amitriptyline-treated mice and tumor growth was monitored once tumors have been established (n= 8-9). (B) CD44 and IFN-γ expression of Tlls were analyzed by flow cytometry. Results from 2 independent experiments are depicted as mean ± SEM. Statistical analysis was performed by 2way ANOVA with Sidak’s multiple comparisons or Student’s t-test. (*p < 0.05, **p < 0.01, ****p < 0.0001).

**Supplemental Fig. S2.**
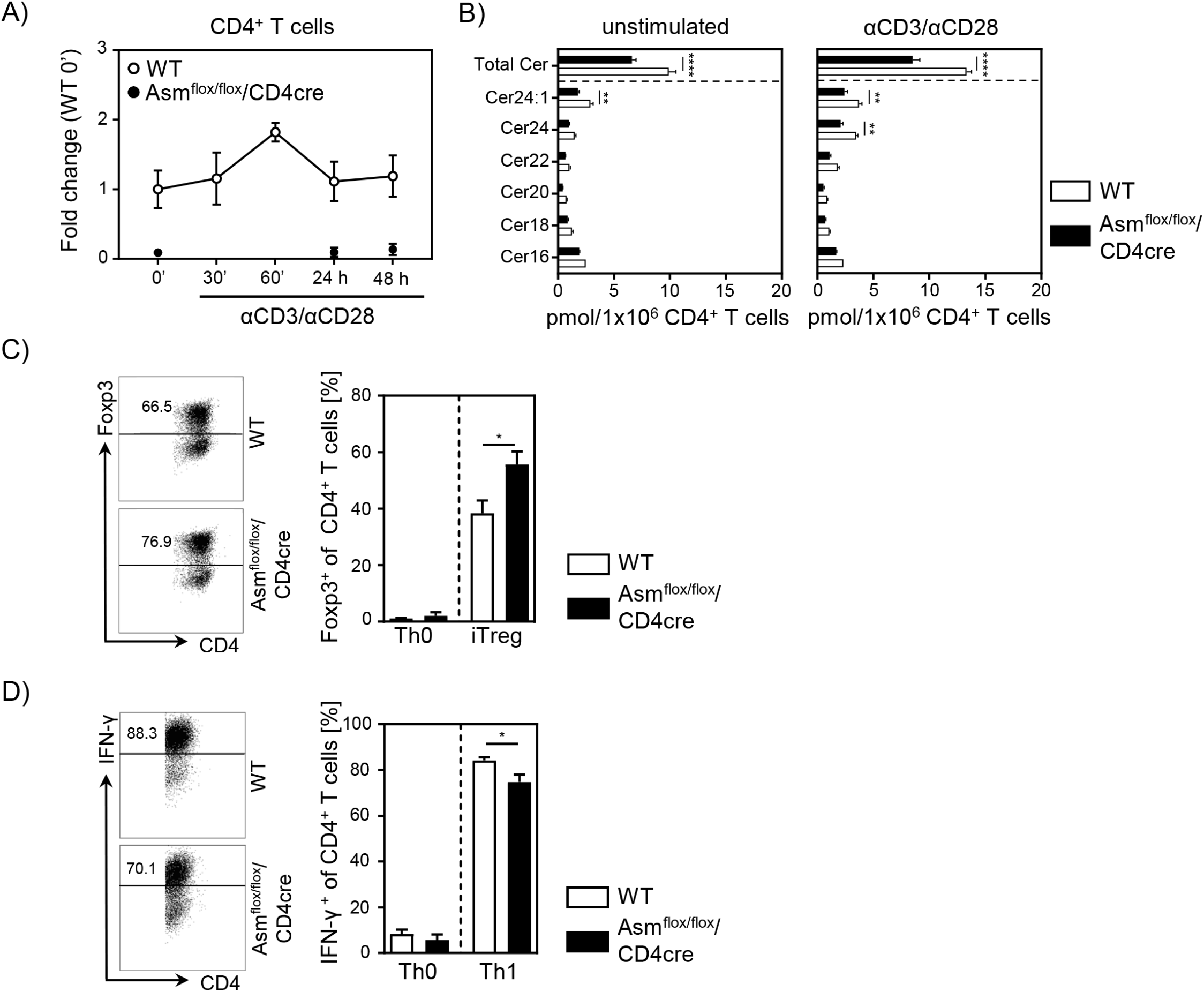
(A) Isolated CD4^+^ T cells from Asm^flox/flox^/CD4cre mice and WT littermates were either left unstimulated or stimulated with anti-CD3 and anti-CD28 antibodies for indicated time points. mRNA expression of Asm *(Smpd1)* was analyzed by RT-qPCR (n= 6-8). (B) Ceramide levels of unstimulated or anti-CD3/anti-CD28 stimulated (24 h) CD4^+^ T cells from Asm^flox/flox^/CD4cre mice and WT littermates were determined by mass spectrometry (n= 4). (C) Sorted CD4^+^CD25- T cells from Asm^flox/flox^/CD4cre and WT mice were stimulated with anti-CD3 and anti-CD28 antibodies in the presence of IL-2 and TGF-131 (iTreg). Respective controls (Th0) were only stimulated with anti-CD3 and anti-CD28 antibodies. After 3 days, Treg differentiation was analyzed by Foxp3 expression (n= 6-7). Representative dot plots are depicted in the left panel. (D) In order to differentiate naïve CD4^+^CD25- T cells into Th1 cells, isolated cells were cultured in the presence of anti CD3, anti-CD28, anti-lL-4, and IL-12 (Th1), or only stimulated with anti-CD3 and anti-CD28 (Th0). After 6 days, Th1 differentiation was assessed by IFN-γ expression (n= 5-6). Representative dot plots are shown in the left panel. Results from 2 to 3 independent experiments are depicted as mean ± SEM. Statistical analysis was performed by 2way ANOVA with Sidak’s multiple comparisons or Student’s t-test. (*p < 0.05, **p < 0.01, ****p < 0.0001).

**Supplemental Fig. S3.**
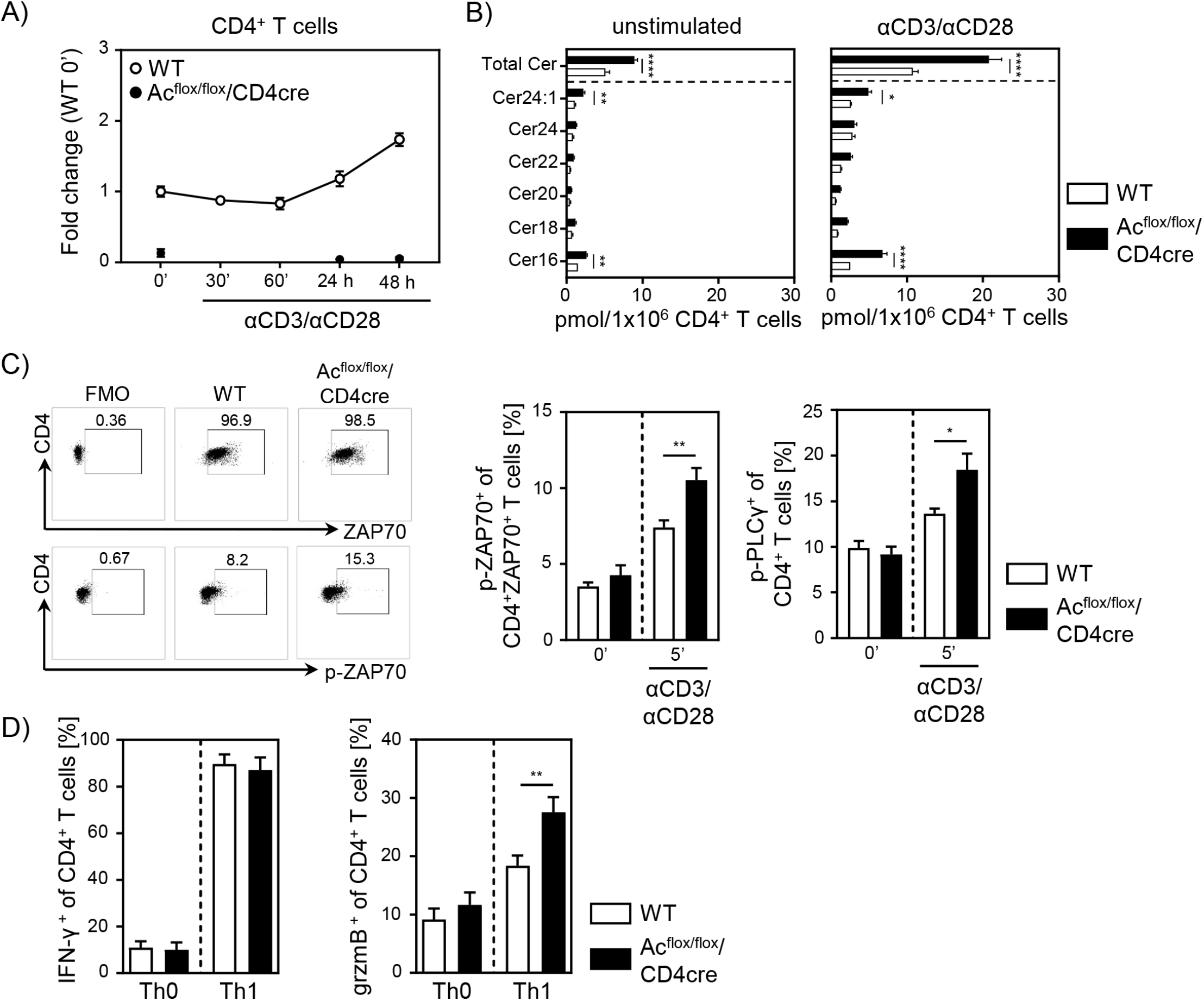
(A) Isolated CD4^+^ T cells from Ac^flox/flox^/CD4cre and WT mice were either left unstimulated or stimulated with anti-CD3 and anti-CD28 antibodies for indicated time points. mRNA expression of Ac *(Asah1)* was analyzed by RT-qPCR (n= 3-4). (B) Ceramide levels of unstimulated and anti-CD3/anti-CD28 stimulated (24 h) CD4^+^ T cells from Ac^flox/flox^/CD4cre and WT mice were determined by mass spectrometry (n= 4). (C) For TCR signaling analysis, splenocytes from Ac^flox/flox^/CD4cre and WT mice were left unstimulated (0’) or stimulated with anti-CD3 and anti-CD28 antibodies for 5 (5’) min. Afterwards, samples were analyzed for phospho ZAP70 of gated ZAP70^+^CD4^+^ T cells and phospho-PLCy of gated CD4^+^ T cells by flow cytometry (n= 4-7). Representative dot plots and FMOs are shown in the left panel. (D) In order to differentiate na”fve CD4^+^ CD25- T cells from Ac^flox/flox^/CD4cre and WT mice into Th1 cells, isolated cells were cultured in the presence of anti-CD3, anti-CD28, anti-lL-4, and IL-12 (Th1), or only stimulated with anti-CD3 and anti-CD28 antibodies (Th0). After 6 days, cells were analysed for IFN-γ and granzyme B expression by flow cytometry (n= 9). Results from 2 to 3 independent experiments are depicted as mean ± SEM. Statistical analysis was performed by 2way ANOVA with Sidak’s multiple comparisons or Student’s t-test. (*p < 0.05, **p < 0.01, ****p < 0.0001).

